# Nasal postbiotic therapy restores NALT architecture and enhances respiratory innate immunity in protein-malnourished mice

**DOI:** 10.64898/2026.04.08.717191

**Authors:** Ivir Maximiliano, Vasile Brenda, Gutiérrez Florencia, Alvarez Villamil Eduardo, Alvarez Susana, Salva Susana

## Abstract

**Background:** Malnutrition compromises mucosal immunity, especially in the respiratory tract, increasing susceptibility to pathogens like *Streptococcus pneumoniae*. This study assessed whether nasal administration of *Lacticaseibacillus rhamnosus* CRL1505 or its peptidoglycan could promote the recovery of nasopharynx-associated lymphoid tissue (NALT) structure and functionality, thereby enhancing resistance to *S. pneumoniae* infection in protein-malnourished mice.

**Methods:** Male Swiss albino mice were fed to a protein-free diet to induce malnutrition, followed by nutritional repletion with or without nasal supplementation of CRL1505 or its peptidoglycan. Resistance to *S. pneumoniae* infection, NALT architecture, immune cell composition in NALT and regional lymph nodes, and nasal cytokine production were evaluated.

**Results:** Protein deficiency caused marked NALT atrophy, immune cell depletion, and heightened susceptibility to *S. pneumoniae*. Nutritional repletion alone partially reversed these effects. In contrast, nasal supplementation with CRL1505 or its postbiotic fully restored NALT structure and cellularity, normalized lymphoid and myeloid populations, and reduced pathogen burden. Both treatments increased B and T lymphocytes, immature B cells, dendritic cells, and macrophages. The postbiotic also enhanced MHCII expression and balanced neutrophil-like Gr-1⁺ cells. Notably, immune enhancement was evident even before infection, indicating a mucosal priming effect. Cytokine levels in nasal fluids remained largely unchanged.

**Conclusions:** Nasal delivery of *L. rhamnosus* CRL1505 or its postbiotic effectively reestablished NALT integrity and mucosal immunity in malnourished mice, providing significant protection against respiratory pathogens. These findings support the development of nasal immunobiotic formulations as non-invasive interventions to bolster respiratory defenses in immunocompromised hosts.

## Introduction

The nasal mucosa is essential for immunosurveillance of inhaled respiratory pathogens. In mice, the nasopharynx-associated lymphoid tissue (NALT) is the only organized mucosa-associated lymphoid structure in the upper respiratory tract and functions similarly to Peyer’s patches in the gut and Waldeyer’s ring in humans^1–3^. NALT consists of organized lymphoid aggregates located beneath the respiratory epithelium of the nasal cavity and serves as an inductive site for mucosal immune responses^4^. Despite its strategic anatomical location and functional relevance, NALT remains comparatively less characterized than other mucosa-associated lymphoid tissues, particularly under conditions of nutritional stress. However, the impact of protein malnutrition on the structural and cellular organization of NALT, as well as on its capacity to sustain local immune responses, has been insufficiently explored.

Malnutrition, particularly in developing countries, is a leading cause of secondary immunodeficiency, leading to atrophy of mucosa-associated lymphoid tissues^5^, and increased susceptibility to *Streptococcus pneumoniae* due to impaired innate responses^6^.

In this context, immunobiotic lactic acid bacteria and their postbiotics have emerged as safe, non-invasive strategies for restoring respiratory immunity in malnourished hosts. Nasal administration of *Lacticaseibacillus rhamnosus* CRL1505 enhances immune responses^7–9^, and its peptidoglycan retains immunomodulatory activity even without bacterial viability^10, 11^.

Despite these findings, the present study evaluates the immunoregulatory effects of nasal administration of *L. rhamnosus* CRL1505 and its postbiotic on the structural organization and innate immune profile of NALT in protein-malnourished mice. By focusing on this undercharacterized mucosal inductive site, this work aims to expand current knowledge of upper respiratory mucosal immunity and to provide experimental support for the development of nasal immunointerventions in immunocompromised populations.

## Materials and methods

### Microorganisms

*Lacticaseibacillus rhamnosus* CRL1505, a well-characterized immunobiotic strain from the CERELA culture collection (*San Miguel de Tucumán, Argentina*), was cultured in Man-Rogosa-Sharpe (MRS) broth following rehydration in a nutrient medium (1.5% peptone, 1% tryptone, 0.5% meat extract, pH 7). After 12 hours of incubation at 37 °C, the bacteria were harvested by centrifugation, washed under sterile conditions, and resuspended in sterile phosphate-buffered saline (PBS, 0.01 mol/L, pH 7.2)^7,10–13^.

*Streptococcus pneumoniae* serotype 6B, isolated from a human respiratory sample (*Malbrán Institute, Buenos Aires*), was cultured in Todd Hewitt broth (THB) for 18 hours at 37 °C and washed in sterile PBS^14^.

### Postbiotic preparation

Peptidoglycan was extracted from CRL1505 using a modified version of Shida et al.^15,10^. The resulting preparation was confirmed to be free of lipoteichoic acid, wall teichoic acid, and nucleic acid, with phosphorus levels below the detection limit (<10 nmol/mg). Endotoxin levels were <0.01 EU/mg, as assessed using a chromogenic LAL assay (Thermo Scientific, USA). Fourier-transform infrared spectroscopy confirmed its identity as lactic acid bacterial peptidoglycan.

### Animals and feeding regimen

Three-week-old male Swiss albino mice were obtained from CERELA animal facility. Malnutrition was induced by feeding a protein-free diet (PFD) for 21 days^16^. Malnourished mice (weighing ∼50 ± 3% of well-nourished controls) were then repleted with a balanced conventional diet (BCD) and randomly assigned to three groups (Fig. 1): 1) BCD group: BCD for 7 days; 2) BCD+Lr05 group: BCD for 7 days plus nasal CRL1505 (10^8^ CFU/mouse/day) during the final 2 days. 3) BCD+PG05 group: BCD for 7 days plus nasal CRL1505 peptidoglycan (8 µg/ml) during the final 2 days^10^. Malnourished control mice maintained on PFD (MNC). A fifth group of well-nourished mice (WNC) received BCD *ad libitum*. Diet compositions have been previously described^16^. Two independent experiments were conducted using nine animals per group.

**Figure 1.**
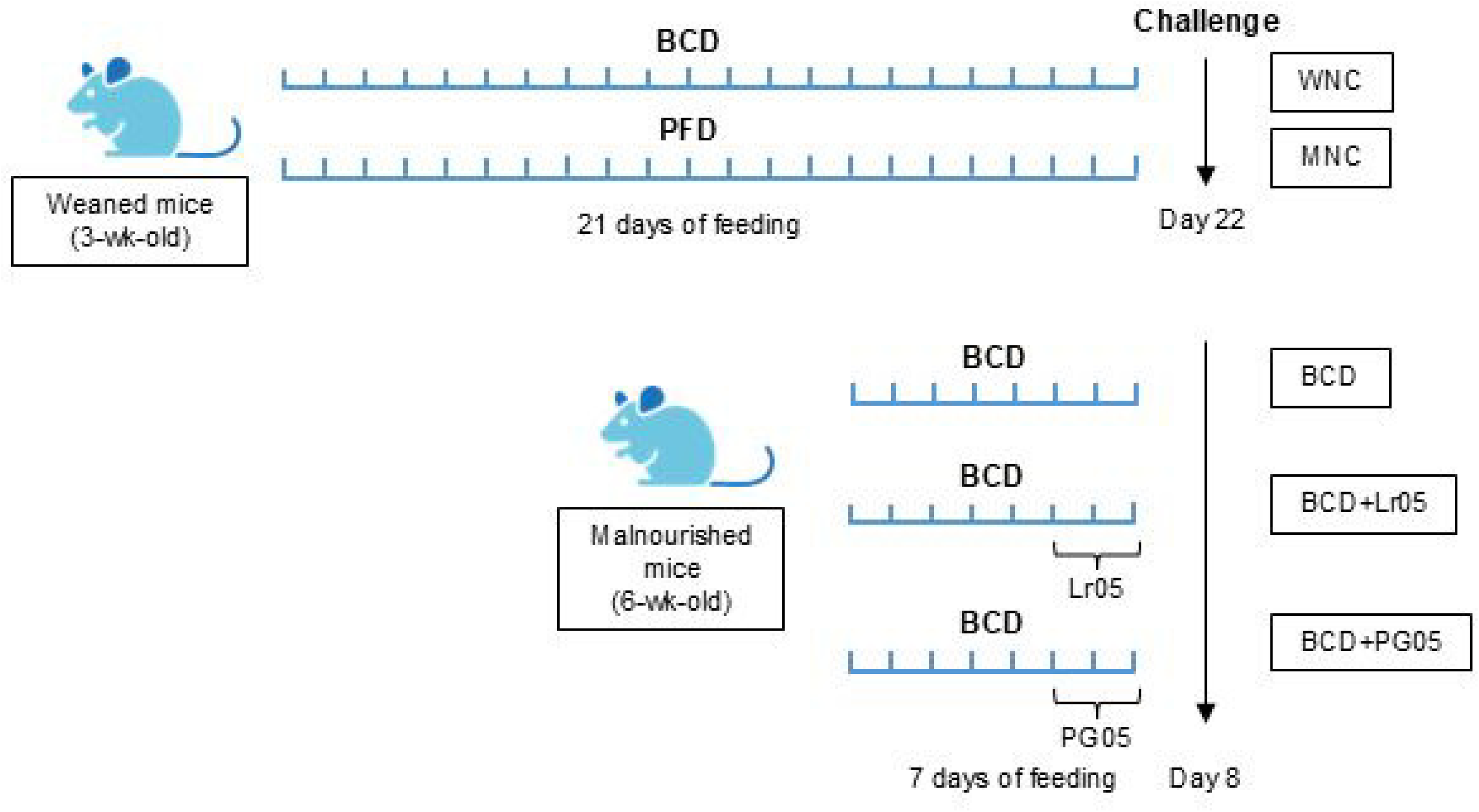
Schematic representation of the malnutrition model and experimental design. The model illustrates the induction of malnutrition in three-week-old Swiss albino mice through a protein-free diet (PFD) for 21 days. Following the malnutrition phase, malnourished mice were repleted with a balanced conventional diet (BCD) and randomly assigned to three groups: (1) BCD group, repleted with BCD for 7 days; (2) BCD+Lr05 group, repleted with BCD for 7 days and receiving intranasal administration of *Lacticaseibacillus rhamnosus* CRL1505 (10⁸ CFU/mouse/day) during the final 2 days; and (3) BCD+PG05 group, repleted with BCD for 7 days and receiving intranasal administration of CRL1505-derived peptidoglycan (8 µg/ml) during the final 2 days. A malnourished control group (MNC) was maintained on PFD without treatment, while a well-nourished control group (WNC) received BCD *ad libitum* throughout the study. After repletion, mice were challenged with *Streptococcus pneumoniae* to assess immune responses. The diagram outlines the timeline for diet administration, treatments, and sample collection. Lr05: *Lacticaseibacillus rhamnosus* CRL1505. PG05: CRL1505-derived peptidoglycan.

The selected dose (10⁸ CFU/mouse/day) and peptidoglycan concentration (8 µg/ml) were based on previous studies demonstrating effective modulation of respiratory innate immunity after nasal administration without inducing adverse inflammatory effects^7–10^. The 2-day administration schedule was chosen according to earlier reports showing that short-term nasal stimulation is sufficient to induce measurable changes in mucosal immune cell populations and cytokine production in the upper respiratory tract^7–9^. The intervention was applied during the final phase of nutritional repletion to evaluate its impact on early immune restoration.

Animal protocols were approved by the Institutional Committee for the Care and Use of Laboratory Animals of CERELA, Tucumán, Argentina (protocols CRL-CICUAL-IBT-2024/2A and CRL-CICUAL-IBT-2024/7A), and followed national and international guidelines.

### Histological analysis of NALT

Post-treatments, nasal cavities were collected, flushed with PBS, and fixed in 4% formalin. After dehydrated and paraffin embedding, sections (4 μm) were stained with hematoxylin-eosin. All samples were coded and assessed blindly under light microscopy.

### Microbiological analysis of nasal lavage

Nasal lavage was performed by inserting a catheter via the trachea and flushing with 400 µL PBS containing 1% heparin. Recovered fluids were plated on Brain Heart Infusion (BHI) blood agar, MacConkey agar and MRS agar, and incubated at 37°C for 24-48 hours. Results were expressed as CFU/mL.

### Pneumococcal infection

After feeding treatment, mice were intranasally challenged with 25 µL of *S. pneumoniae* (10⁷ CFU in PBS) per nostril^10^. To facilitate alveoli delivery, mice were held upright for 2 minutes post-instillation. Euthanasia was performed on day 0 (pre-challenge) or day 2 post-infection.

### Isolation of cells from lymphoid tissues

On days 0 and 2 post-infection, cervical and axillary lymph nodes (CLN and ALN) and NALT were collected. Lymph nodes were dissected following the protocol of Van den Broeck et al.^17^ and homogenized in RPMI 1640 medium supplemented with 2% fetal bovine serum (FBS). NALT was extracted following Asanuma and Heritage’s methods^18,19^. The upper palate was surgically exposed and removed to extract NALT aggregates, which were then flushed with RPMI-FBS solution. Cell viability was assessed using trypan blue exclusion.

### Flow cytometry

Cells were incubated with Fc Block (anti-CD32/CD16) prior to staining with monoclonal antibodies (BD PharMingen) targeting: CD3, CD4, CD8, B220, CD24, IgM, CD11b, CD11c, MHCII, Gr-1, F4/80, and CD103. Biotinylated antibodies were visualized using streptavidin-PercP. Data were acquired using a BD FACSCalibur™ flow cytometer and analyzed with FlowJo software (TreeStar). Absolute cell counts were calculated by multiplying the total number viable cells by the percentage of each cell population.

### Cytokine quantification

Cytokine levels (TNF-α, IFN-γ, IL-10) were measured in nasal lavage fluid on day 2 post-infection using ELISA kits (eBioscience, USA). Values were expressed in pg/mL based on standard curves.

### Statistical analysis

Data were presented as mean ± SD from triplicate experiments and analyzed using GraphPad Prism 6.0. After confirming normal distribution, two-way ANOVA was performed, followed by Tukey’s post hoc test for group comparisons. A *p*-value < 0.05 was considered statistically significant.

## Results

### L. rhamnosus CRL1505 and its postbiotic restore NALT structure and cellular composition during infection after protein deprivation

Previous studies showed that nasal administration of *L. rhamnosus* CRL1505 or its postbiotic reduced *S. pneumoniae* load in the lungs and blood^10^. Building on this, we investigated their effects on NALT under protein-deficient conditions. in the WNC mice, NALT appeared as typical lymphoid aggregates in the inferior turbinates (Fig. 2A), while malnourished mice showed pronounced NALT atrophy, reduced body weight, and lower NALT cell counts. Although all dietary repletion protocols partially improved these alterations (Fig. 2B), only BCD+Lr05 and BCD+PG05 significantly increased NALT cell numbers (Fig. 2C) and restored the NALT-to-body weight ratio (Fig. 2D). Mice receiving BCD alone maintained severely affected.

**Figure 2.**
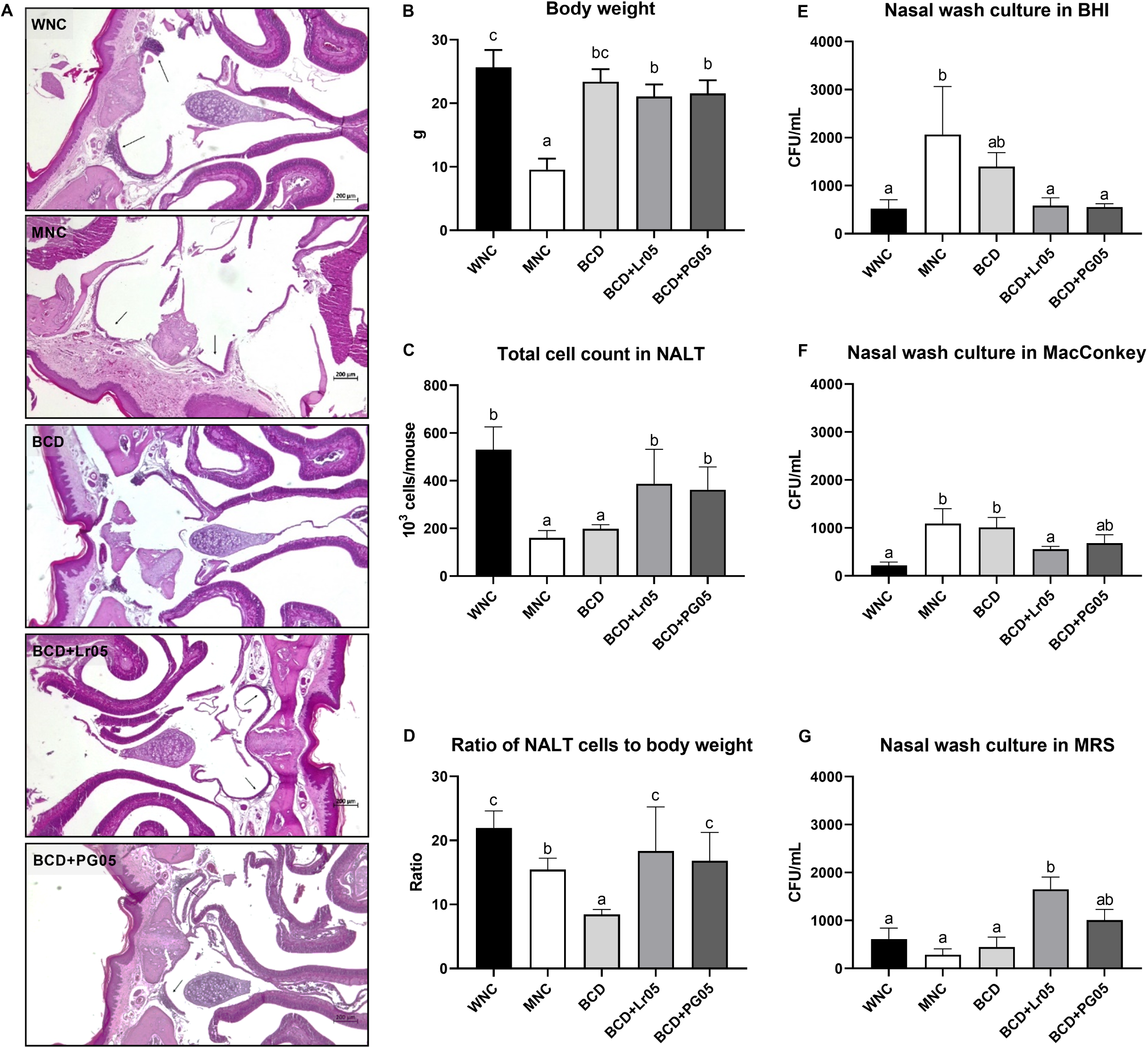
Effect of *L. rhamnosus* CRL1505 and its postbiotic on NALT recovery. Malnourished mice were repleted with a balanced conventional diet (BCD) and randomly assigned to three groups: (1) BCD group, repleted with BCD for 7 days; (2) BCD+Lr05 group, repleted with BCD for 7 days and receiving intranasal administration of *Lacticaseibacillus rhamnosus* CRL1505 during the final 2 days; and (3) BCD+PG05 group, repleted with BCD for 7 days and receiving intranasal administration of CRL1505-derived peptidoglycan during the final 2 days. Malnourished (MNC) and well-nourished (WNC) mice were used as controls. (A) NALT samples were collected, fixed, stained with hematoxylin and eosin, and examined under a light microscope (400× magnification). Arrows indicate NALT. (B) Body weight. (C) Total cell count in NALT. (D) Ratio of NALT cells to body weight. Nasal lavage fluid was collected and plated on BHI agar (E), MacConkey agar (F), and MRS agar (G) to evaluate bacterial counts. The results represent data from two independent experiments, with nine animals per group. Results are expressed as mean ± standard deviation. ^a, b, c^ Means in a bar with different letters (a < b < c) were significantly different (*p* < 0.05). Lr05: *Lacticaseibacillus rhamnosus* CRL1505. PG05: CRL1505-derived peptidoglycan. NALT: nasal-associated lymphoid tissue. BHI: brain heart infusion. MRS: de Man–Rogosa–Sharpe.

NALT atrophy correlated with increased nasal bacterial load, as shown by higher colony counts on BHI and MacConkey agar (Fig. 2E, F). This was normalized only in groups treated nasally with CRL1505 or its postbiotic (Fig. 2E, F). Moreover, only the BCD+Lr05 group showed a significant rise in beneficial bacterial colonies on MRS agar (Fig. 2G), suggesting selective enrichment of protective microbiota.

### L. rhamnosus CRL1505 and its postbiotic modulate the innate immune response in NALT

To explore the mechanisms of protection, we analyzed NALT immune cell populations, focusing on innate response. Protein malnutrition led to significant reductions in total lymphocyte, total B cells (B220⁺), mature B cells (B220^High^CD24^Low^), total T cells (CD3^+^), and CD4⁺/ CD8⁺ subsets (Fig. 3A-C, E-G) prior to infection. Nasal supplementation with *L. rhamnosus* CRL1505 or its postbiotic restored these populations, while BCD alone had no effect. Notably, only the nasal treatments increased immature B cells (B220^Low^CD24^High^), suggesting stimulation of local lymphopoiesis (Fig. 3D).

**Figure 3.**
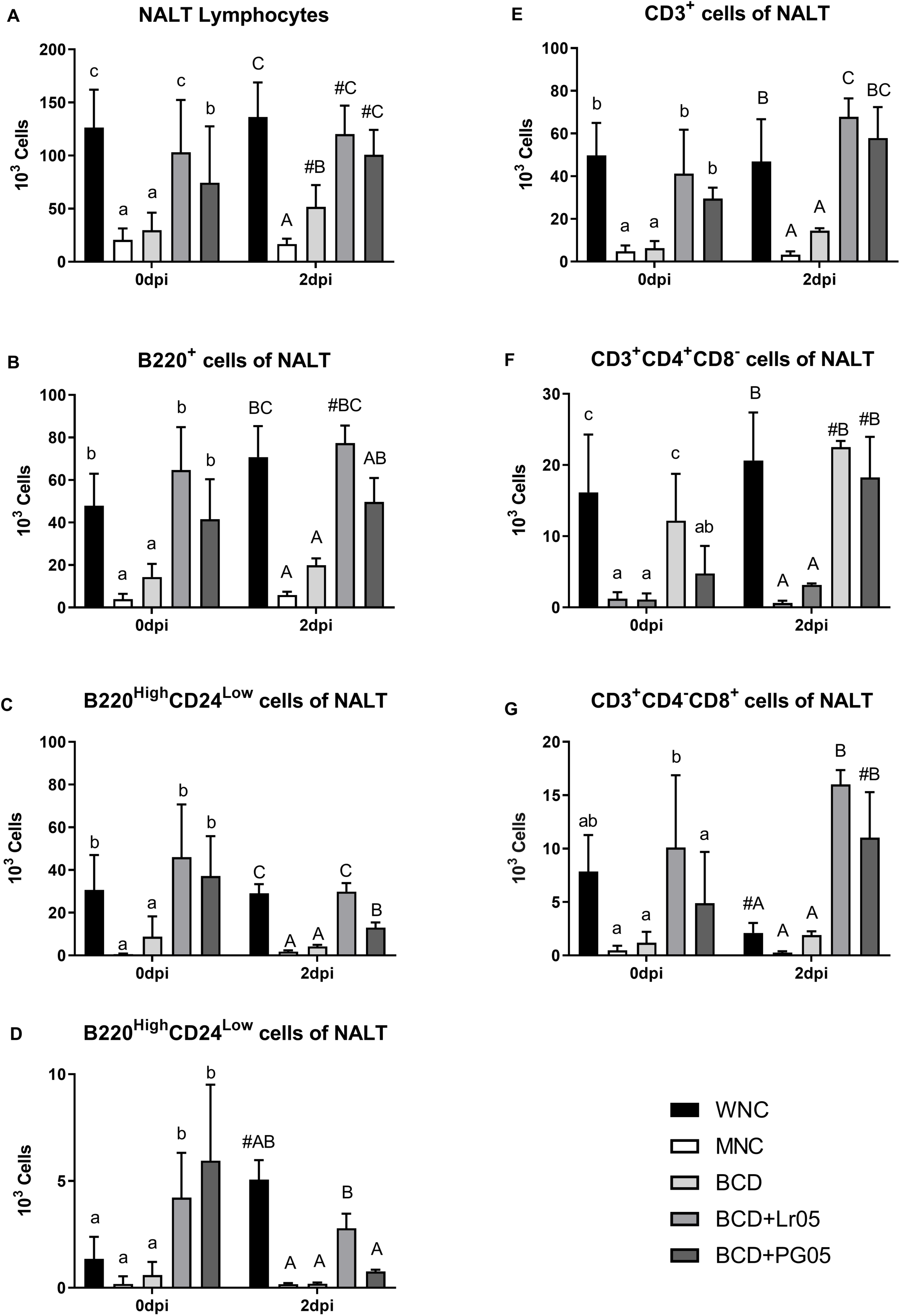
Recovery of NALT lymphocytes. Malnourished mice were repleted with a balanced conventional diet (BCD) and randomly assigned to three groups: (1) BCD group, repleted with BCD for 7 days; (2) BCD+Lr05 group, repleted with BCD for 7 days and receiving intranasal administration of *Lacticaseibacillus rhamnosus* CRL1505 during the final 2 days; and (3) BCD+PG05 group, repleted with BCD for 7 days and receiving intranasal administration of CRL1505-derived peptidoglycan during the final 2 days. Malnourished (MNC) and well-nourished (WNC) mice were used as controls. (A) Number of total lymphocytes (FFSC vs SSC). (B) Number of total B cells (B220^+^ cells). (C) Number of mature B cells (B220^High^CD24^Low^ cells). (D) Number of immature B cells (B220^Low^CD24^High^ cells). (E) Number of total T cells (CD3^+^ cells). (F) Number of T helper cells (CD3^+^CD4^+^CD8^-^ cells). (G) Number of T cytotoxic cells (CD3^+^CD4^-^CD8^+^ cells). The results represent data from two independent experiments. Nine animals from each group were used. Results are expressed as mean ± standard deviation. a,b,c Means in a bar with different letters (a < b < c) were significantly different (*P* < 0.05). Capital letters are used for day 2 post-infection. # Means significant difference with day 0. Lr05: *Lacticaseibacillus rhamnosus* CRL1505. PG05: CRL1505-derived peptidoglycan. NALT: nasal-associated lymphoid tissue.

In contrast, BCD+Lr05 and BCD+PG05 groups exhibited lymphocyte levels comparable to or higher than WNC mice, including enhanced total lymphocytes, B cells, T cells, and CD4⁺ T cells.

Following infection, the WNC group showed only minor shifts in lymphocyte profiles, whereas MNC and BCD groups remained severely immunosuppressed (Fig. 3). In contrast, BCD+Lr05 and BCD+PG05 groups exhibited lymphocyte levels comparable to or higher than WNC mice, including enhanced total lymphocytes, B cells, T cells, and CD4⁺ T cells (Fig. 3).

Protein malnutrition reduced CD11b⁺CD11c⁻ cells and CD103⁺ dendritic cells in NALT (Fig. 4A, B). Nasal supplementation with *L. rhamnosus* CRL1505 or its postbiotic, unlike the BCD group, restored these populations and significantly increased F4/80⁺ macrophages (Fig. 4C). Post-infection, CD11b⁺CD11c⁻ cells declined in well-nourished mice but remained unchanged in malnourished ones. In contrast, CD103⁺ and F4/80⁺ cells were further reduced by infection in malnourished animals (Fig. 4B, C). Both nasal treatments increased all three myeloid populations post-infection, whereas BCD alone had no effect (Fig. 4A-C). Consistently, before infection, only the BCD+Lr05 group showed enhanced MHCII expression, as indicated by increased mean fluorescence intensity (MFI). After infection, MHCII expression declined in most groups, except BCD+PG05, which maintained the highest levels (Fig. 4D). In malnourished mice, the percentage of Gr-1⁺ cells in NALT increased (Fig. 4E), although all groups showed a reduction in their absolute numbers (Fig. 4F). Infection induced an increase in both the percentage and absolute number of these cells in the WNC group. Importantly, only the BCD+Lr05 and BCD+PG05 groups showed a significant increase in the percentage of Gr-1⁺ cells and successfully restored their absolute numbers in NALT (Fig. 4E, F).

**Figure 4.**
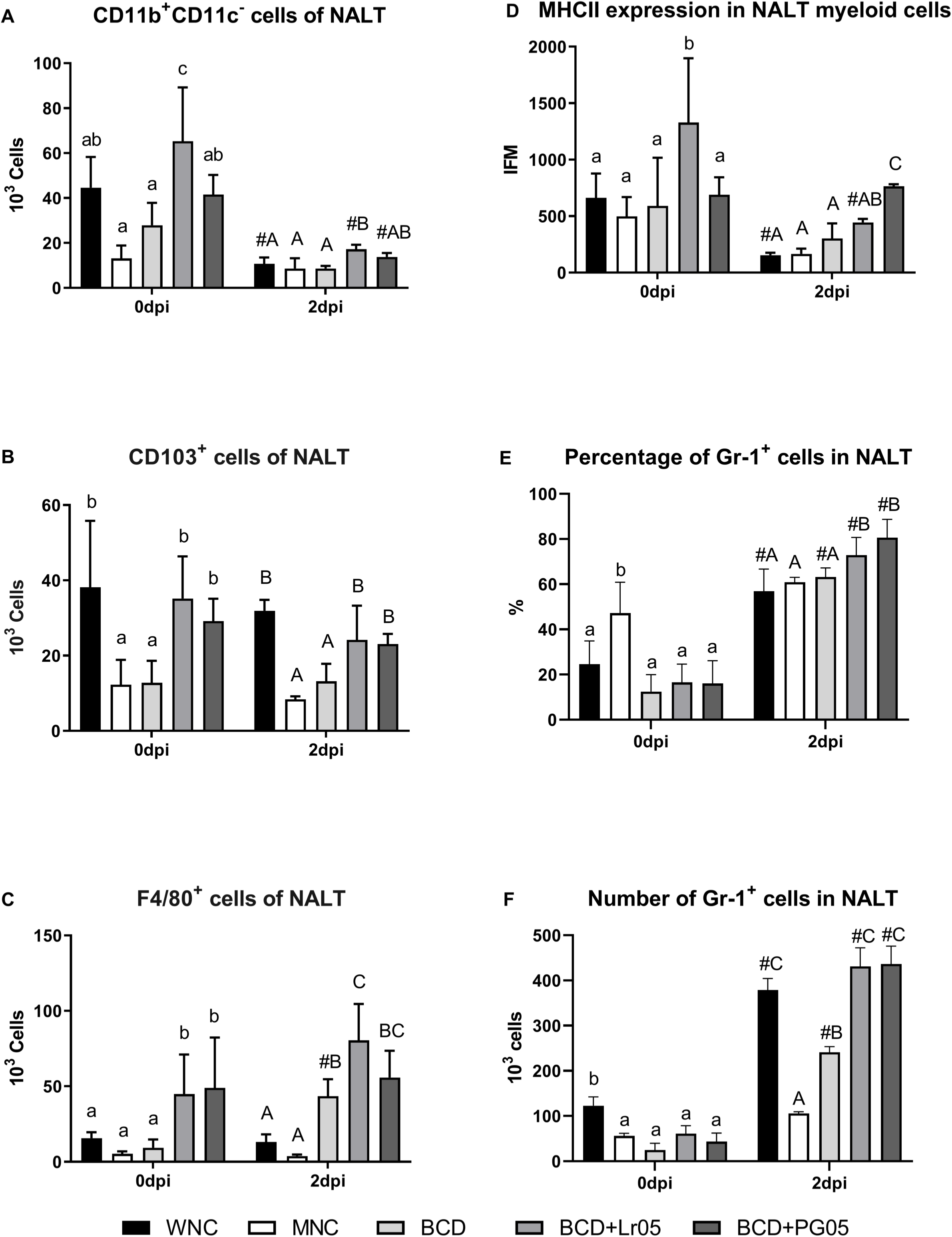
Recovery of NALT antigen-presenting cells. Malnourished mice were repleted with a balanced conventional diet (BCD) and randomly assigned to three groups: (1) BCD group, repleted with BCD for 7 days; (2) BCD+Lr05 group, repleted with BCD for 7 days and receiving intranasal administration of *Lacticaseibacillus rhamnosus* CRL1505 during the final 2 days; and (3) BCD+PG05 group, repleted with BCD for 7 days and receiving intranasal administration of CRL1505-derived peptidoglycan during the final 2 days. Malnourished (MNC) and well-nourished (WNC) mice were used as controls. (A) Number of CD11b^+^CD11c^-^ cells. (B) Number of CD103^+^ cells. (C) Number of F4/80+ cells. (D) Mean fluorescence intensity of MHCII. (E) Percentage of Gr-1 cells. (F) Number of Gr-1^+^ cells. The results represent data from two independent experiments. Nine animals from each group were used. Results are expressed as mean ± standard deviation. a,b,c Means in a bar with different letters (a < b < c) were significantly different (*p* < 0.05). Capital letters are used for day 2 post-infection. # Means significant difference with day 0. Lr05: *Lacticaseibacillus rhamnosus* CRL1505. PG05: CRL1505-derived peptidoglycan. NALT: nasal-associated lymphoid tissue.

To further characterize the local immune response, TNF-α, IL-10, and IFN-γ levels were measured in nasal lavage fluid. Malnutrition did not significantly alter these cytokines, though slight increases in TNF-α and IFN-γ were observed in the BCD and BCD+Lr05 groups, respectively (Fig. 5).

**Figure 5.**
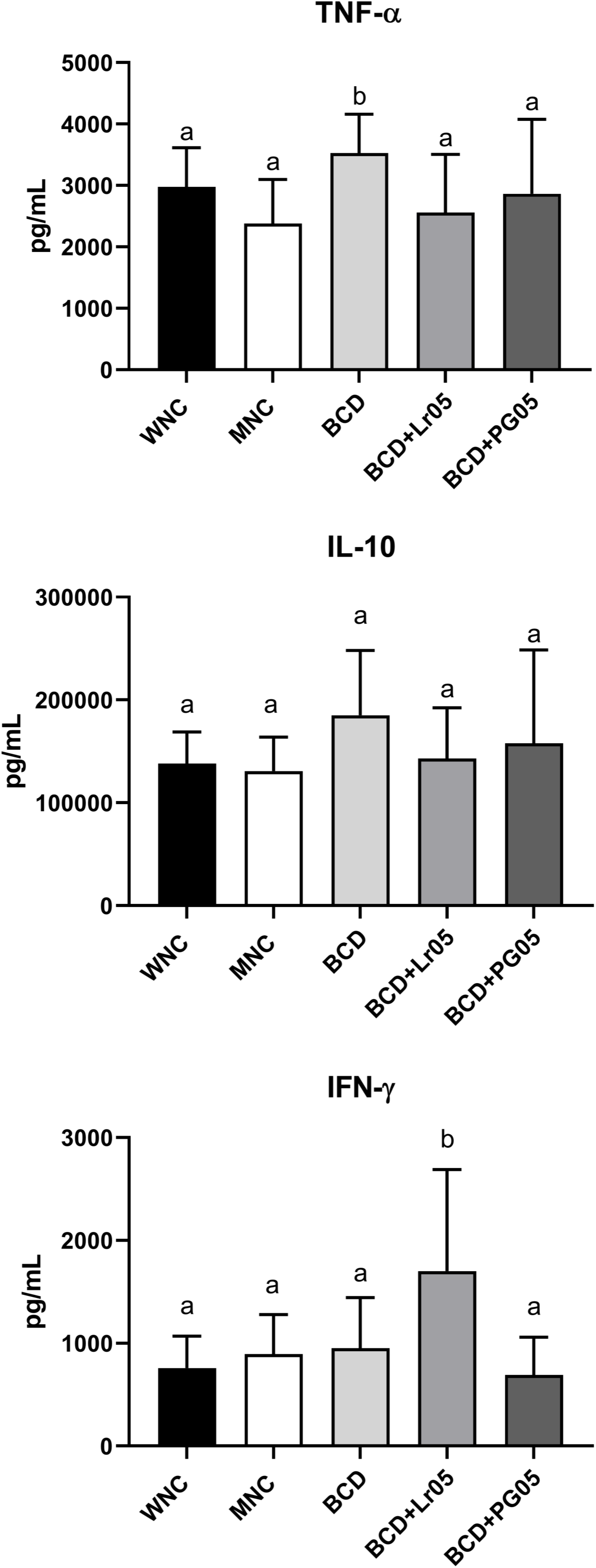
NALF levels of cytokines. Malnourished mice were repleted with a balanced conventional diet (BCD) and randomly assigned to three groups: (1) BCD group, repleted with BCD for 7 days; (2) BCD+Lr05 group, repleted with BCD for 7 days and receiving intranasal administration of *Lacticaseibacillus rhamnosus* CRL1505 during the final 2 days; and (3) BCD+PG05 group, repleted with BCD for 7 days and receiving intranasal administration of CRL1505-derived peptidoglycan during the final 2 days. Malnourished (MNC) and well-nourished (WNC) mice were used as controls. TNF-α, IL-10 and INF-γ were expressed in picograms (pg)/ml using the standard curve performed with different concentrations of the corresponding cytokine. The results represent data from two independent experiments. Nine animals from each group were used. Results are expressed as mean ± standard deviation. a,b Means in a bar with different letters (a < b) were significantly different (*p* < 0.05). Lr05: *Lacticaseibacillus rhamnosus* CRL1505. PG05: CRL1505-derived peptidoglycan. NALF: nasal lavage fluid.

### L. rhamnosus CRL1505 and its postbiotic reverse infection-associated alterations in lymph nodes induced by protein deprivation

Given the crucial role of cervical lymph nodes in regional immunity, we examined structural and cellular changes following infection and dietary treatments. Malnutrition significantly reduced all analyzed cell populations in cervical lymph nodes (Fig. 6). While dietary repletion partially restored lymphocyte and B cell numbers, only nasal supplementation with L. rhamnosus CRL1505 or its postbiotic fully normalized lymphocytes, total B cells, total T cells, and CD8⁺ T cells (Fig. 6A, B, E, G). Additionally, these nasal treatments increased immature B cells and CD4⁺ T cells beyond levels seen in the BCD group (Fig. 6C, F). Infectious decreased total lymphocytes only in the WNC group’s cervical nodes (Fig. 6A). Notably, nasal-treated groups exhibited higher counts across all cell populations compared to WNC mice post-infection (Fig. 6).

**Figure 6.**
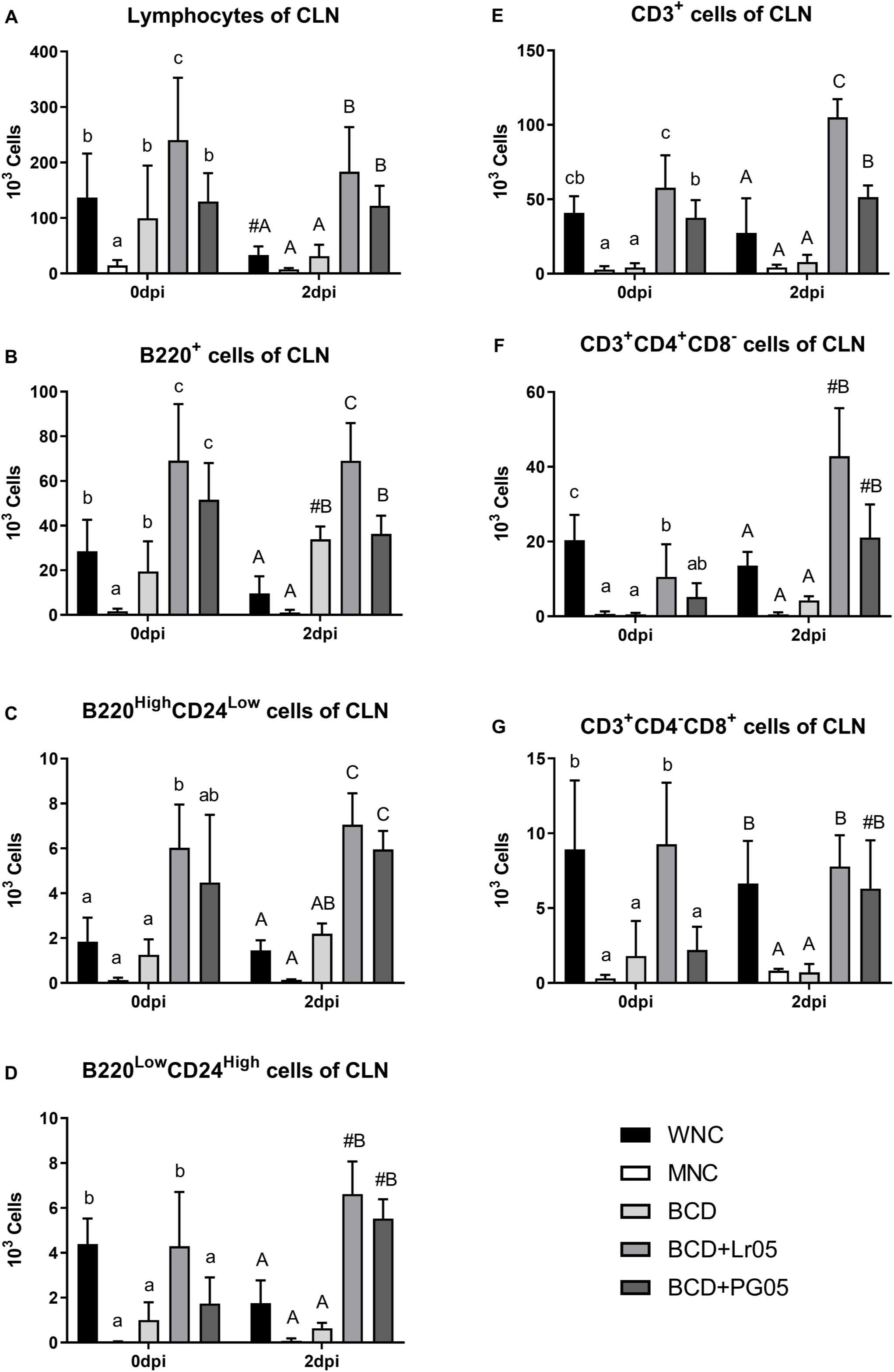
Recovery of CLN cells. Malnourished mice were repleted with a balanced conventional diet (BCD) and randomly assigned to three groups: (1) BCD group, repleted with BCD for 7 days; (2) BCD+Lr05 group, repleted with BCD for 7 days and receiving intranasal administration of *Lacticaseibacillus rhamnosus* CRL1505 during the final 2 days; and (3) BCD+PG05 group, repleted with BCD for 7 days and receiving intranasal administration of CRL1505-derived peptidoglycan during the final 2 days. Malnourished (MNC) and well-nourished (WNC) mice were used as controls. (A) Number of total lymphocytes (FFSC vs SSC). (B) Number of total B cells (B220^+^ cells). (C) Number of mature B cells (B220^High^CD24^Low^ cells). (D) Number of immature B cells (B220^Low^CD24^High^ cells). (E) Number of total T cells (CD3^+^ cells). (F) Number of T helper cells (CD3^+^CD4^+^CD8^-^ cells). (G) Number of T cytotoxic cells (CD3^+^CD4^-^CD8^+^ cells). The results represent data from two independent experiments. Nine animals from each group were used. Results are expressed as mean ± standard deviation. a,b,c Means in a bar with different letters (a < b < c) were significantly different (*p* < 0.05). Capital letters are used for day 2 post-infection. # Means significant difference with day 0. Lr05: *Lacticaseibacillus rhamnosus* CRL1505. PG05: CRL1505-derived peptidoglycan. CLN: cervical lymph node.

To assess whether immunomodulation extended beyond local nodes, we evaluated axillary lymph nodes. Malnutrition reduced all lymphocyte populations studied, and BCD alone failed to reverse these deficits (Fig. 7). Nasal supplementation with *L. rhamnosus* CRL1505 restored the total lymphocytes, total and mature B cells, total T cells, and CD4⁺ T cells to normal levels (Fig. 7A, B, C, E, F), and increased immature B cells beyond those of WNC group (Fig. 7G). Peptidoglycan treatment also improved total, mature, and immature B cells, as well as total T cells (Fig. 7B, C, D, E). Infection in WNC group increased total lymphocytes, total T cells, and CD8⁺ T cells (Fig. 7A, E, G). Post-infection, the MNC and BCD groups displayed lower cell counts than WNC group, while nasal treatments partially restored these populations (Fig. 7).

**Figure 7.**
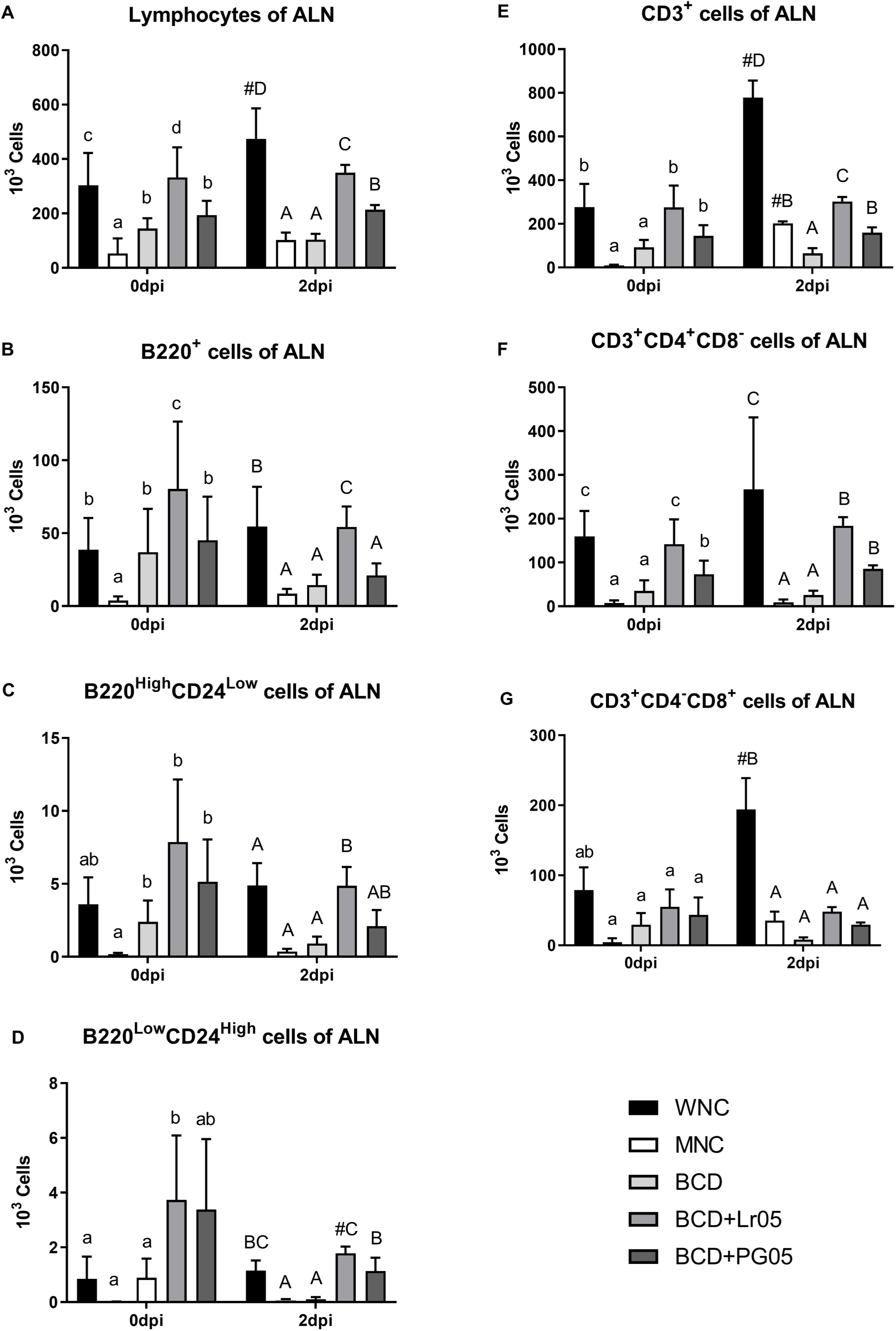
Recovery of ALN cells. Malnourished mice were repleted with a balanced conventional diet (BCD) and randomly assigned to three groups: (1) BCD group, repleted with BCD for 7 days; (2) BCD+Lr05 group, repleted with BCD for 7 days and receiving intranasal administration of *Lacticaseibacillus rhamnosus* CRL1505 during the final 2 days; and (3) BCD+PG05 group, repleted with BCD for 7 days and receiving intranasal administration of CRL1505-derived peptidoglycan during the final 2 days. Malnourished (MNC) and well-nourished (WNC) mice were used as controls. (A) Number of total lymphocytes (FFSC vs SSC). (B) Number of total B cells (B220^+^ cells). (C) Number of mature B cells (B220^High^CD24^Low^ cells). (D) Number of immature B cells (B220^Low^CD24^High^ cells). (E) Number of total T cells (CD3^+^ cells). (F) Number of T helper cells (CD3^+^CD4^+^CD8^-^ cells). (G) Number of T cytotoxic cells (CD3^+^CD4^-^CD8^+^ cells). The results represent data from two independent experiments. Nine animals from each group were used. Results are expressed as mean ± standard deviation. a,b,c,d Means in a bar with different letters (a < b < c < d) were significantly different (*p* < 0.05). Capital letters are used for day 2 post-infection. # Means significant difference with day 0. Lr05: *Lacticaseibacillus rhamnosus* CRL1505. PG05: CRL1505-derived peptidoglycan. ALN: axillary lymph node.

## Discussion

This study is the first to demonstrate that intranasal administration of *L. rhamnosus* CRL1505 or its postbiotic (peptidoglycan) promotes the structural and immunological recovery of NALT in a mouse model of protein malnutrition. By specifically focusing on this underexplored upper respiratory inductive site, our findings provide direct experimental evidence that targeted nasal immunomodulation can reverse malnutrition-associated alterations at the level of organized lymphoid tissue. These findings highlight the potential of this strategy to improve resistance against *S. pneumoniae* in immunocompromised hosts.

A key finding was that both the probiotic and its peptidoglycan restored the NALT architecture disrupted by malnutrition. Importantly, this study characterizes for the first time the structural and cellular alterations of NALT under protein-deficient conditions and demonstrates their reversibility through nasal intervention. This recovery correlated with increased total lymphoid cells and normalized T and B cell profiles, suggesting reactivation of local lymphopoiesis. In contrast, dietary repletion alone did not reverse these immune impairments, indicating that nutritional recovery per se is insufficient to reestablish nasal mucosal immune competence. These findings are consistent with previous reports of systemic and pulmonary immune restoration by CRL1505^7,8,10,11,20^, although few studies outside this group have explored its nasal effects^21,22^.

Consistent with histological and immunophenotypic findings, CRL1505 or PG05 treatment reduced bacterial load in nasal lavage, supporting the role of enhanced local innate immunity in protecting against respiratory pathogens. Additionally, both treatments enriched beneficial nasal microbiota, as indicated by increased MRS agar colony counts, suggesting that immune restoration was accompanied by improved microbial balance at the mucosal surface^23^. In our study, the restoration of myeloid populations (CD11b⁺, CD103⁺, F4/80⁺), recovery of MHC II expression, and increased Gr-1⁺ cells point to coordinated activation of innate immune components, enhancing mucosal surveillance and response. These coordinated cellular changes provide a functional framework linking NALT reorganization with improved pathogen control. These changes contrast with the dysbiosis-linked immune imbalance observed in conditions like chronic rhinosinusitis and adenoid hypertrophy^24,25^.

*S. pneumoniae* is an opportunistic pathogen that initially colonizes the nasopharynx. While its prevalence is low in healthy adults, it can reach up to 65% in children, posing a great risk for malnourished individuals^26,27^. In our model, malnourished mice exhibited a severely impaired immune response, characterized by reduced myeloid populations (particularly macrophages and dendritic cells), and low MHCII expression. Macrophage depletion in NALT compromised early immune responses^28^. Although a neutrophil proportion increased, their absolute numbers remained low, indicating inadequate recruitment. Dendritic cell loss further weakened innate-adaptive crosstalk. This immune dysfunction led to elevated bacterial loads in the lungs and bloodstream, reflecting a diminished ability to control infection.

Mice renourished with BCD exhibited a disease progression similar to that of malnourished mice, reinforcing the concept that anthropometric recovery does not necessarily parallel mucosal immune restoration^11,29^. In contrast, nasal treatments with *L. rhamnosus* CRL1505 or its postbiotic produced strong immunomodulatory effects, restoring total cellularity, improving myeloid and lymphoid populations, and enhancing antigen-presenting cell activation. Notably, although both interventions were effective, the live probiotic induced stronger responses in macrophage recovery and IFN-γ production, indicating differential immunomodulatory capacities rather than complete equivalence between treatments.

Treatments also restored lymphocyte populations in NALT and draining lymph nodes, a key step for effective early adaptive responses closely linked to innate immunity^30^. *L. rhamnosus* CRL1505 proved more effective than its postbiotic in recovering and activating T lymphocytes, supporting its broader stimulation of the mucosa–lymph node axis. These immunological improvements were associated with educed bacterial dissemination to the lungs and bloodstream, supporting a direct relationship between NALT immune reconstitution and enhanced infection control. The postbiotic also conferred notable beneficials, including CD103⁺cell restoration, increased MHCII expression, and reduction pathogen burden.

## Conclusions

Overall, *L. rhamnosus* CRL1505 and its postbiotic emerge as promising candidates for non-invasive immunonutritional therapies targeting vulnerable populations. This study provides experimental evidence that localized nasal immunomodulation can reestablish NALT organization and improve early antimicrobial defense in the context of protein malnutrition. The effectiveness of the postbiotic highlights its potential as a safe alternative when live bacteria are unsuitable, although the differential magnitude of certain immune parameters suggests that live and non-viable preparations are not fully interchangeable.

Future studies should further characterize the cellular and investigate the molecular pathways involved in NALT reactivation and evaluate long-term efficacy in models or chronic infection to support translational development.

## CRediT authorship contribution statement

The authors’ responsibilities were as follows: SS, SA, and MI designed the research; MI, BV, FG, EVA and SS conducted the research; MI and SS analyzed the data; and SS, MI and SA wrote the paper. SS had primary responsibility for final content. All authors read and approved the final manuscript.

## Acknowledgements

This work was supported in part by grants from the Consejo Nacional de Investigaciones Científicas y Técnicas (CONICET) (PIP 0545) and the Agencia Nacional de Promoción Científica y Tecnológica (ANPCyT) (PICT 2021-0278).

## References

1. Zuercher AW, Coffin SE, Thurnheer MC, Fundova P, Cebra JJ. Nasal-associated lymphoid tissue is a mucosal inductive site for virus-specific humoral and cellular immune responses. J Immunol. 2002;168:1796–1803. 10.4049/jimmunol.168.4.1796.

2. Kiyono H, Fukuyama S. NALT-versus Peyer’s-patch-mediated mucosal immunity. Nat Rev Immunol. 2004;4:699–710. 10.1038/nri1439.

3. Rodríguez-Monroy MA, Rojas-Hernández S, Moreno-Fierros L. Phenotypic and functional differences between lymphocytes from NALT and nasal passages of mice. Scan J Immunol. 2007:65;276–288. 10.1111/j.1365-3083.2006.01898.x.

4. Sepahi A, Salinas I. The evolution of nasal immune systems in vertebrates. Mol Immunol. 2016:69;131–138. 10.1016/j.molimm.2015.09.008.

5. Savino W, Durães J, Maldonado-Galdeano C, Perdigón G, Mendes-da-Cruz DA, Cuervo P. Thymus, undernutrition, and infection: Approaching cellular and molecular interactions. Front Nutr. 2022;9: 948488, 10.3389/fnut.2022.948488.

6. Marangu D, Zar HJ. Childhood pneumonia in low-and-middle-income countries: An update. Paediatr Respir Rev. 2019;32:3–9. 10.1016/j.prrv.2019.06.001.

7. Barbieri N, Villena J, Herrera M, Salva S, Alvarez S. Nasally administered *Lactobacillus rhamnosus* accelerate the recovery of humoral immunity in B lymphocyte-deficient malnourished mice. J Nutr 2013;143:227–235. 10.3945/jn.112.165811.

8. Barbieri N, Herrera M, Salva S, Villena J, Alvarez S. *Lactobacillus rhamnosus* CRL1505 nasal administration improves recovery of T-cell mediated immunity against pneumococcal infection in malnourished mice. Benef Microbes. 2017;8:393–405. 10.3920/BM2016.0152.

9. Herrera M, Salva S, Villena J, Barbieri N, Marranzino G, Alvarez S. Dietary supplementation with Lactobacilli improves emergency granulopoiesis in protein-malnourished mice and enhances respiratory innate immune response. PLoS One. 2014;9(4):e90227. 10.1371/journal.pone.0090227.

10. Kolling Y, Salva S, Villena J, Marranzino G, Alvarez S. Non-viable immunobiotic *Lactobacillus rhamnosus* CRL1505 and its peptidoglycan improve systemic and respiratory innate immune response during recovery of immunocompromised-malnourished mice. Int Immunopharmacol. 2015;25:474–484. 10.1016/j.intimp.2015.02.006.

11. Kolling Y, Salva S, Villena J, Alvarez S. Are the immunomodulatory properties of *Lactobacillus rhamnosus* CRL1505 peptidoglycan common for all Lactobacilli during respiratory infection in malnourished mice? PLoS One. 2018;13(3):e0194034. 10.1371/journal.pone.0194034.

12. Salva S, Villena J, Alvarez S. Immunomodulatory activity of *Lactobacillus rhamnosus* strains isolated from goat milk: impact on intestinal and respiratory infections. Int J Food Microbiol. 2011;141:82–89. 10.1016/j.ijfoodmicro.2010.03.013.

13. Salva S, Alvarez S. The role of microbiota and immunobiotics in granulopoiesis of immunocompromised hosts. Front Immunol. 2017;8:507. 10.3389/fimmu.2017.00507.

14. Villena J, Racedo S, Agüero G, Bru E, Medina M, Alvarez S. *Lactobacillus casei* improves resistance to pneumococcal respiratory infection in malnourished mice. J Nutr. 2005;135:1462–1469. 10.1093/jn/135.6.1462.

15. Shida K, Kiyoshima-Shibata J, Kaji R, Nagaoka M, Nanno M. Peptidoglycan from lactobacilli inhibits interleukin-12 production by macrophages induced by *Lactobacillus casei* through Toll-like receptor 2-dependent and independent mechanisms. Immunology. 2009;128:e858–69. 10.1111/j.1365-2567.2009.03095.x.

16. Salva S, Villena J, Racedo S, Alvarez S, Agüero G. *Lactobacillus casei* addition to a repletion diet-induced early normalization of cytokine profils during a pneumococcal infection in malnourished mice. Food Agric Immunol. 2008;19:195–211. 10.1080/09540100802247243.

17. Van den Broeck W, Derore A, Simoens P. Anatomy and nomenclature of murine lymph nodes: descriptive study and nomenclatory standardization in BALB/cAnNCrl mice. J Immunol Methods. 2006;312:12–19. 10.1016/j.jim.2006.01.022.

18. Asanuma H, Thompson AH, Iwasaki T, et al. Isolation and characterization of mouse nasal-associated lymphoid tissue. J Immunol Methods. 1997;202:123–131. 10.1016/s0022-1759(96)00243-8.

19. Heritage PL, Underdown BJ, Arsenault AL, Snider DP, McDermott MR. Comparison of murine nasal-associated lymphoid tissue and Peyer’s patches. Am J Respir Crit Care Med. 1997;156:1256–1262. 10.1164/ajrccm.156.4.97-03017.

20. Barbieri N, Salva S, Herrera M, Villena J, Alvarez S. Nasal priming with *Lactobacillus rhamnosus* CRL1505 stimulates mononuclear phagocytes of immunocompromised malnourished mice: Improvement of respiratory immune response. Probiotics Antimicrob Proteins. 2020;12:494–504. 10.1007/s12602-019-09551-8.

21. Vintiñi EO, Medina M. Immune response in nasopharynx, lung, and blood elicited by experimental nasal pneumococcal vaccines containing live or heat-killed lactobacilli as mucosal adjuvants. Can J Physiol Pharmacol. 2014;92:124–131. 10.1139/cjpp-2013-0227.

22. Makino T, Yamashita M, Takeuchi N, Kabuki T, Hattori M, Yoshida T. Lactobacillus helveticus SBT2171 alleviates allergic symptoms in a murine model for pollen allergy. Biosc Biotechnol Biochem. 2019;83:2298–2306. 10.1080/09168451.2019.1654847.

23. Costalonga M, Cleary PP, Fischer LA, Zhao Z. Intranasal bacteria induce Th1 but not Treg or Th2. Mucosal Immunol. 2009;2:85–95. 10.1038/mi.2008.67.

24. Liu W, Jiang H, Liu X, et al. Altered intestinal microbiota enhances adenoid hypertrophy by disrupting the immune balance. Front Immunol. 2023;14:1277351. 10.3389/fimmu.2023.1277351.

25. Jain R, Waldvogel-Thurlow S, Darveau R, Douglas R. Differences in the paranasal sinuses between germ-free and pathogen-free mice. Int Forum Allergy Rhinol. 2016;6:631–637. 10.1002/alr.21712

26. Weiser JN, Ferreira DM, Paton JC. *Streptococcus pneumoniae*: transmission, colonization and invasion. Nat Rev Microbiol. 2018;16:355–367. 10.1038/s41579-018-0001-8.

27. Ibrahim MK, Zambruni M, Melby CL, Melby PC. Impact of childhood malnutrition on host defense and infection. Clin Microbiol Rev. 2017;30:919–971. 10.1128/cmr.00119-16.

28. Aberdein JD, Cole J, Bewley MA, Marriott HM, Dockrell DH. Alveolar macrophages in pulmonary host defence – the unrecognized role of apoptosis as a mechanism of intracellular bacterial killing. Clin Exp Immunol. 2013;174:193–202. 10.1111/cei.12170.

29. Rytter MJH, Kolte L, Briend A, Friis H, Christensen VB. The immune system in children with malnutrition-A systematic review. PLoS One. 2014,9:e105017. 10.1371/journal.pone.0105017.

30. Mettelman RC, Allen EK, Thomas PG. Mucosal immune responses to infection and vaccination in the respiratory tract. Immunity. 2022;55:749–780. 10.1016/j.immuni.2022.04.013.

